# Understanding the General Principles of T Cell Engagement by Multiscale Computational Simulations

**DOI:** 10.1101/2023.06.07.544116

**Authors:** Zhaoqian Su, Steven C. Almo, Yinghao Wu

## Abstract

The use of bispecific antibodies as T cell engagers can bypass the normal TCR-MHC interaction, redirect the cytotoxic activity of T-cells, and lead to highly efficient tumor cell killing. However, this immunotherapy also causes significant on-target off-tumor toxicologic effects, especially when they were used to treat solid tumors. In order to avoid these adverse events, it is necessary to understand the fundamental mechanisms during the physical process of T cell engagement. We developed a multiscale computational framework to reach this goal. The framework combines simulations on the intercellular and multicellular levels. On the intercellular level, we simulated the spatial-temporal dynamics of three-body interactions among bispecific antibodies, CD3 and TAA. The derived number of intercellular bonds formed between CD3 and TAA were further transferred into the multicellular simulations as the input parameter of adhesive density between cells. Through the simulations under various molecular and cellular conditions, we were able to gain new insights of how to adopt the most appropriate strategy to maximize the drug efficacy and avoid the off-target effect. For instance, we discovered that the low antibody binding affinity resulted in the formation of large clusters at the cell-cell interface, which could be important to control the downstream signaling pathways. We also tested different molecular architectures of the bispecific antibody and suggested the existence of an optimal length in regulating the T cell engagement. Overall, the current multiscale simulations serve as a prove-of-concept study to help the future design of new biological therapeutics.

**SIGNIFICANCE:** T-cell engagers are a class of anti-cancer drugs that can directly kill tumor cells by bringing T cells next to them. However, current treatments using T-cell engagers can cause serious side-effects. In order to reduce these effects, it is necessary to understand how T cells and tumor cells interact together through the connection of T-cell engagers. Unfortunately, this process is not well studied due to the limitations in current experimental techniques. We developed computational models on two different scales to simulate the physical process of T cell engagement. Our simulation results provide new insights into the general properties of T cell engagers. The new simulation methods can therefore serve as a useful tool to design novel antibodies for cancer immunotherapy.

## Introduction

New cancer treatments that rely on augmenting the patient’s immune system first reached the clinic in the late 1990s as the result of our expanding understanding of adaptive immunity [1–3]. These treatments, known as immunotherapies, enhance the antitumor activities of T cells through a range of approaches [4]. Current T-cell immunotherapies fall into three major categories. Immune checkpoint inhibitors, one of the first immunotherapies, elicit global stimulation of the T cell repertoire, which is associated with enhanced antitumor activity [5, 6]. The second strategy, adoptive T cell therapies, includes chimeric antigen receptor (CAR) T cell therapy, which relies on the harvesting and *in vitro* culturing of CAR T cells and is time-consuming that may be prohibitively costly [7–9]. Lastly, T cell engagers (TCEs), such as bispecific T cell engagers (BiTEs) redirect T cells to cancers using a protein construct that bridges CD3 on T cells and tumor-associated antigens (TAA) [10, 11]. This engagement releases perforin and granzyme, leading to highly efficient tumor cell killing [12]. Over 100 BiTEs are in the clinical trials [13]; however, these biologics can elicit significant on-target off-tumor effects, especially when they were used to treat solid tumors [14]. To reduce these adverse events, it is necessary to understand the fundamental mechanisms that are operative during the physical process of T cell engagement. For instance, how can TCEs be designed to more effectively bind to their target receptors on T cells and tumor cells, and how do different molecular and cellular factors regulate this process? Unfortunately, it is challenging to address these questions on a quantitative level due to current experimental limitations.

Computational simulation is a less time-consuming and labor-intensive approach to reveal mechanistic insight into complex biological systems, and these methods have become a fundamental component of modern drug development [15–17]. The majority of computational drug design studies have concentrated on, and made important contributions to, the discovery of therapeutic antibodies [18, 19], peptide-based biomolecules [20, 21], and small chemical compounds [22]. In contrast, computational analyses of multi-specific biologics have been hampered by their molecular complexity as well as the limitations underlying the most commonly used computational approaches including molecular dynamic (MD) simulations [23–25] and protein-protein docking [26–28]. The enormous computational demands of MD simulations, for example, represents significant hurdles for applying these strategies to explore macromolecular systems like bispecific biologics over biologically relevant timescales. Moreover, effective immunotherapies involve the regulation of interactions between drugs and their targets in multicellular systems, while all computational modeling methods used in current drug discovery focus on the atomic details of molecular interactions. Therefore, new approaches that can link biological processes from the molecular to the cellular level are of high interest. Multiscale modeling is a promising technique that can bridge computational simulations on different scales [29, 30], and has been applied to study cancer biology [31, 32]. However, the application of multiscale simulations in understanding the mechanism of T cell engagement has not been documented.

In this study, we developed a multiscale simulation framework to explore the general determinants of T cell engagers, using a prototypical BiTE is used as a test model. A kinetic Monte-Carlo method was first designed to simulate the spatial-temporal dynamics of the three-body interactions involving bispecific biologics recognizing both CD3 and a tumor associated antigen (TAA), CD3 on a cytotoxic T cell and the TAA on a targeted tumor cell. We demonstrate that the number of intercellular bonds formed between CD3 (T cell) and TAA (tumor cell) behave according to the classic Hook effect, with a concave down dependence on concentration (i.e., increasing at low concentration, achievement of maximal interactions at intermediated concentrations and decreasing as the concentration continued to increase). We found that the peak position is sensitive to the density of cell surface receptors as well as the binding affinity between receptors and antibodies. Of particular interest, our simulation results show that the lower binding affinity can lead to the formation of larger receptor clusters at the cell-cell interface. We also demonstrated sensitivity to linker length in the bispecific reagent. The number of intercellular bonds observed between CD3 and TAA in the above intercellular simulations were further converted into the adhesive density between cells and incorporated into a multicellular model. The Langevin dynamics simulation utilized different values of adhesive density as input parameters to drive the dynamics of a mixed system with T cells, tumor cells and normal cells. The behavior of T cell engagers can therefore be explored through a combination between of intercellular and the multicellular simulations. In summary, our results provide new insights into the general principles of T cell engagers and support new multiscale simulation methods as useful tools to design bispecific/multispecific antibodies for cancer immunotherapy.

## Model and Methods

### The kinetic Monte-Carlo simulation of an inter-cellular system

We first simulated the interactions between the bispecific engagers and their targeted membrane receptors on the surfaces of the T cell and tumor cell. A rigid-body (RB) based model was used to represent the intercellular system. Specifically, the plasma membrane of a T cell is represented by the square surface at the bottom of a three-dimensional simulation box, while CD3 molecules are modeled by cylindrical rigid bodies randomly distributed on its surface (**Figure 1a**). Analogously, the plasma membrane of a tumor cell is represented by the square surface at the top of the simulation box, on which cylindrical rigid bodies representing the TAAs are randomly distributed. A defined number of bispecific antibodies are distributed within the three-dimensional volume between the surfaces of the T cell and the tumor cell. Each antibody contains two functional modules that are connected by a linker, and each functional module is modeled by a spherical rigid body. The heights of both CD3 and TAA cylinders are 8nm and their radius is 4nm; the radius of each spherical rigid body is 4nm. The dimension of the cylindrical and spherical rigid bodies are comparable to those of the actual protein structures. The two modules in the bispecific antibody possess non-identical binding sites, one which recognizes CD3 and the other which engages the TAA target. The binding sites of CD3 and TAA receptor are located at the tops of respective rigid body modules, while the binding sites in the bispecific antibody are placed at the tips of both spherical rigid bodies. Using the above model representation and an initial random configuration, the movements of receptors (constrained to two dimensions) and antibodies (three dimensions), as well as their binding interaction, were simulated by a kinetic Monte-Carlo (KMC) algorithm [33] until the system reached equilibrium.

**Figure 1:**
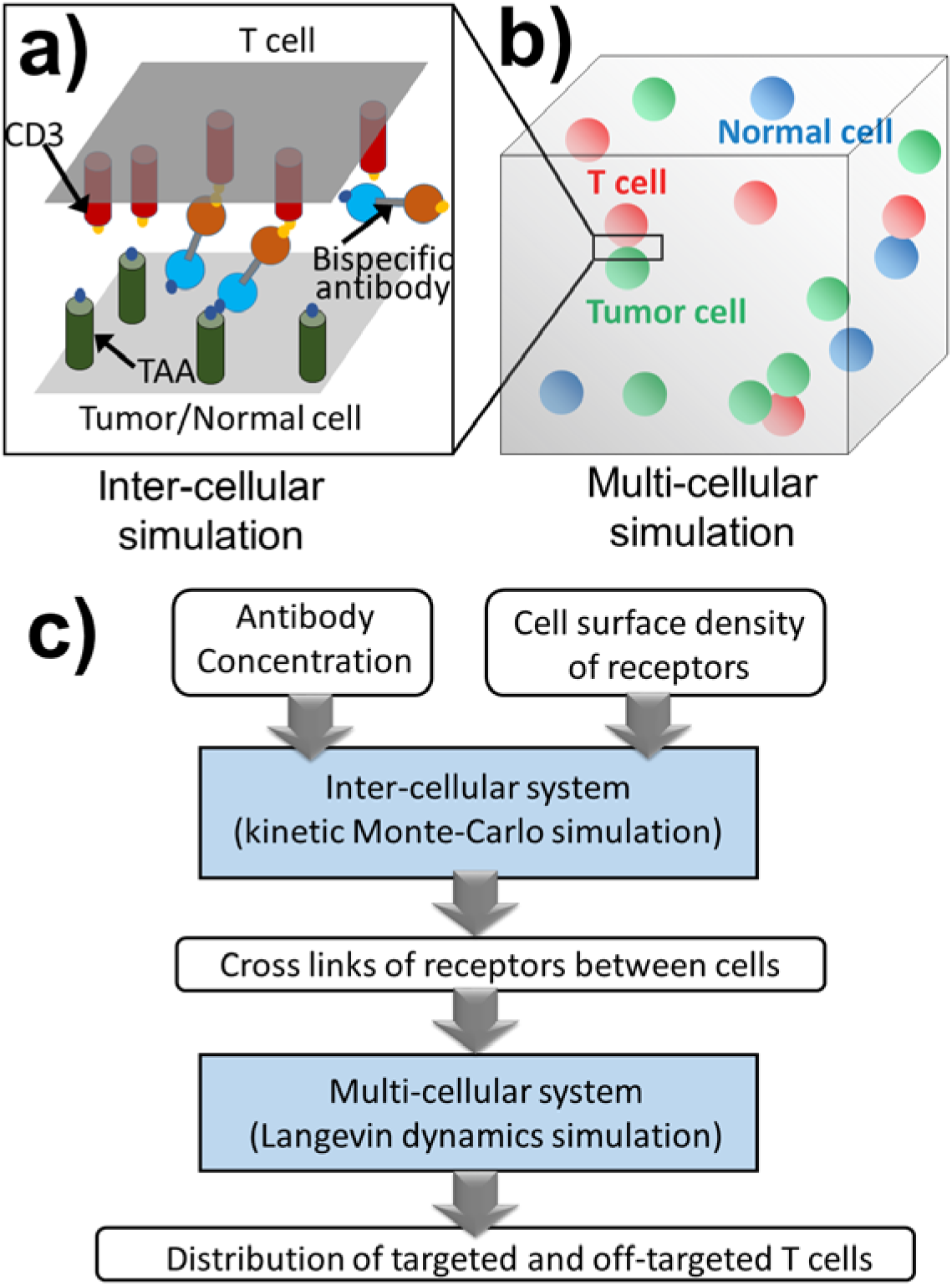
A rigid-body (RB) based kinetic Monte-Carlo simulation was developed to study the binding between the bispecific antibodies and their targeted membrane receptors at the cell-cell interface, which is simplified by two overlapping square planes **(a)**. The derived number of intercellular bonds formed between CD3 and TAA were further transferred into the multicellular simulations, in which numbers of cells are randomly mixed in a cubic simulation box **(b)**. Each one of these cells is denoted by a specific cell type, such as T cell, tumor cell or normal cell. The intercellular and multicellular simulations are integrated into a multiscale computational framework. As shown in **(c)**, the inputs of the framework are surface density of receptors and antibody concentration, and the output is the proportions of tumor cells and normal cells that are targeted by T cells.

Within each simulation step, molecules are randomly selected for stochastic diffusion. Antibodies are free to undergo three-dimensional diffusions throughout the volume of the simulation box. Selected antibodies are translated along a random direction vector and rotated along a random axis, with amplitudes of the movements determined by the defined translational and rotational diffusion constants. Periodic boundary conditions are applied both x and y directions of the simulation box and the antibody is not permitted to move below or below the defined plasma membrane, otherwise, it will be bounced back. In contrast, diffusion of membrane-bound CD3 and TAA are confined within their corresponding two-dimensional plasma membrane. Selected CD3 or TAA molecules are randomly translated on the membrane surface and rotated around an axis normal to the membrane, with the amplitudes of the movements determined by the two-dimensional translational and rotational diffusion constants. Identical 2D periodic boundary conditions are imposed along the x and y directions of the surface. The 3D translation and rotational diffusion constants of a bispecific antibody are 50μm^2^/s and 5°ns^-1^, respectively. The 2D diffusion of receptors is much slower due to the membrane confinement, with the translational and rotational diffusion constants of 10 μm^2^/s and 1°ns^-1^. These values are adopted from our previous studies [34].

Interactions between antibodies and receptors evaluated on the basis of the new configuration generated after the diffusional step. An association between an antibody and its receptor is scored if the distance of their binding sites is below a predetermined cutoff value. The probability to trigger the association is determined by the rate constant *k_on_*. In contrast, the dissociation of a receptor-antibody interaction is determined by the rate constant *k_off_*, in which *k_off_* can be written as *k_off_* = *k_on_* × exp (− Δ*G*_0_ *kT*) where Δ*G_0_* is the binding affinity of the interaction. In order to make the time-scale of our simulations computationally accessible, all *k_on_*s in the system were fixed to a value at the upper bound of a diffusion-limited rate constant, in which association is accelerated by long-range electrostatic interactions [35]. This value corresponds to an experimentally measurable value on the level of 10^9^M^-1^s^-^1. Binding affinities were selected from millimolar (mM) to nanomolar (nM), which is the typical range of interactions between membrane receptors and ligands on the surfaces of immune cells [36].

If one functional module in an antibody binds to its target receptor, the entire receptor-antibody complex will move together on the corresponding plasma membrane, with reduced translational and rotational diffusion constants of 4 μm^2^/s and 0.2°ns^-1^, so that the other vacant module in the antibody is accessible for binding with receptors on the opposite plasma membrane. After an antibody dissociate with its receptor, the two proteins will diffuse separately. In the next simulation step, they will either diffuse farther away from each other or have a chance to re-associate as a geminate recombination if their distance is still below the cutoff. If both modules of an antibody are engaged, one with CD3 and the other with TTA, we assume that the diffusion of an entire ternary complex will be further reduced. For computational simplification, the ternary complex will stop diffusing at the cell-cell interface in current simulation. This diffusion and reaction Monte-Carlo process iterates until the system eventually reaches equilibrium.

### The Langevin dynamics simulation of a multi-cellular system

In addition to the simulation of antibody binding at a single cell-cell interface, we expanded our studies to address the spatial-temporal dynamics of interactions among multiple different cell types. Cells with a given concentration are randomly placed in a cubic simulation box to establish the initial configuration of the system. Each cell type (T cell, tumor cell or normal cell) is modeled as a spherical rigid body with radius of 5μm (**Figure 1b**). Antibodies and cell surface receptors are not explicitly modeled, but their impact on cell-cell interactions is controlled by the adhesive density between cells, as described below. The Langevin dynamics (LD) simulation algorithm is applied to drive the movements and interactions of all cells in the system [37, 38]. The equation of motion used to modify the Cartesian coordinate and velocity of cell *i* has the following form:

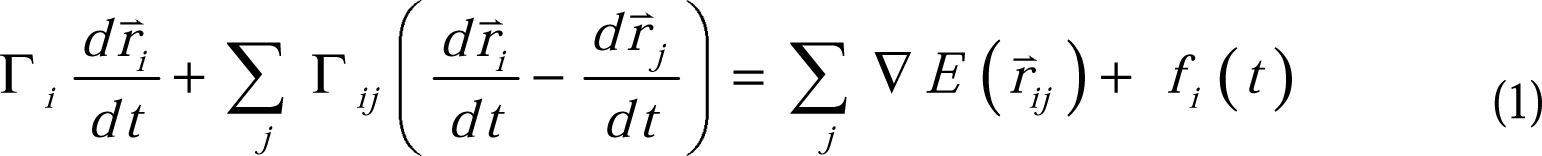

The first term on the left side of equation (1) describes the friction between cell *i dr*^v^ *^k^* and its environment, and 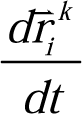 indicates the velocity of the cell. The second term describes *dt* the friction between cell *i* and all its neighboring cells that it contacts. The cell-environments and cell-cell friction tensors are denoted by Γ_i_ and Γ_ij_, respectively. The first term on the right side of equation (1) describes the total forces applied to cell *i*, where *E(r_ij_)* is the potential energy between cell *i* and its neighboring cell *j* and ∇ stands for the gradient operator. The second term on the right side of equation (1) is the stochastic forces that are contributed by the thermal background. More specifically, the potential energy can be broken down into the following two terms.

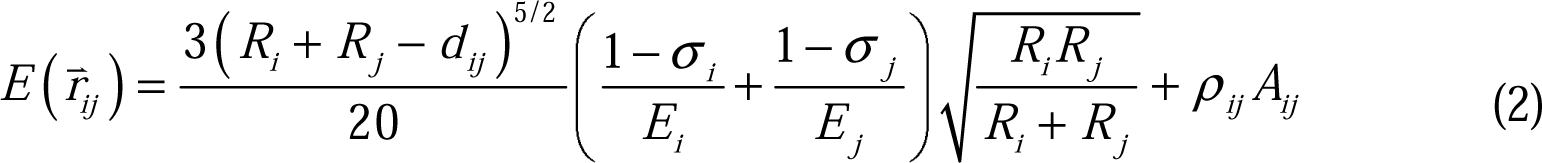

The first term on the right side of equation (2) is adopted from the modified Hertz model and represents the repulsive energy between two cells. The radii of cell *i* and cell *j* are denoted by *R_i_* and *R_j_* and their distance is denoted by *d_ij_*. *E_i_* and σ*_i_*are the elastic moduli and spherical Poisson ratios of the cell, which equals 1kPa and 1/3, respectively. These values are adopted from previous studies [39]. The second term on the right side of equation (2) represents the attractive energy between two cells. *A_ij_* represents the contact area between cells *i* and *j*, and ρ*_ij_* denotes the intercellular adhesive density, which corresponds to the number of ternary complexes formed among bispecific antibodies, CD3 and TAA in the contact area. The value of intercellular adhesion strength depends on the specific types of cells (and their surface receptor density) that form a contact.

Starting from the initial configuration, the force applied to each cell can be calculated by the potential energy given in equation (2) within each step of the Langevin dynamics simulation. Following the force calculation, the velocity and position of all cells in the system are thus updated by the equation of motion described in equation (1). As a result, a new configuration is generated. After repeating this process iteratively, the dynamics of the multicellular system can be evolved until it reaches final equilibrium.

### The framework that integrates the inter-cellular and multi-cellular simulations

Given the kinetic Monte-Carlo and Langevin dynamics simulation algorithms, a multiscale framework was constructed that allows the results from the intercellular system to be incorporated into the multicellular system. We first define the simulation condition, including the surface density of CD3 on T cells, the density of TAAs on the surfaces of tumor cells or normal cells, as well as the concentration of the antibodies. As illustrated in **Figure 1c**, these surface density of receptors and antibody concentration are then used as input parameters for the intercellular simulation. After the intercellular simulation, the number of bonds formed between CD3 and TAA through their interactions with bispecific antibodies can be derived at the interface between T cells and tumor cells, and between T cells and normal cells.

The numbers of intercellular bonds formed at the interface of a given area are, in turn, used as input parameters of the multicellular simulation to determine the adhesion strengths between different cell types. Finally, the energy of attraction between T cells and tumor cells, or between T cells and normal cells, can be calculated by the adhesion strength per surface area (can you put the appropriate units here?). Together with the repulsive energies, the attractive energies are used to calculate forces between cells and guide their movements in the Langevin dynamics simulations. After the multicellular simulations, we can estimate information such as how many tumor cells and normal cells are targeted by T cells by giving their initial densities and ratio in the system. Overall, this multiscale simulation framework enables the exploration of different combinations of antibody concentrations and receptor densities, and the translation of these outcomes to interrogate multicellular systems. Consequently, this approach begins to a provide quantitative and mechanistic understanding of the molecular and cellular factors responsible for the clinical efficacy of multispecific T cell engagers.

## Results

We first applied intercellular simulation to examine how bispecific antibodies regulate the coupling between a T cell and a tumor cell. As described in the Methods, the interface between T cell and tumor cell is modeled as two flat surfaces in close proximity of each other. The size of each square surface is 500 nm along the x and y directions. At the beginning of the simulation, the separation between the two surfaces was set to 70nm (**Figure 2a**), a distance that precludes a bispecific antibody from crosslinking CD3 and TAA on opposing surfaces. Varying numbers of CD3 and TAAs, shown as red and green cylinders in the figure, were randomly distributed on the top and bottom surfaces, respectively. The bispecific antibodies, with the individual affinity modules represented by orange and blue spheres and connected by a 5nm gray linker, were distributed in the volume between the two surfaces. In order to simulate the process during which two cells encounter each other, the intercellular distance was gradually reduced from 70nm to 35nm along the simulation. This value is close to experimentally reported intercellular distances [40]. As a result, CD3, TAA and bispecific antibodies can form ternary complexes at the cell-cell interface, as shown in **Figure 2b**. We evaluated the system with different numbers of receptors on both surfaces, increasing from 10 to 100. This corresponds to the surface density of ∼10^2^ molecules per μm^2^, which was within the range of experimental observation for T cells [41]. After fixing the surface density of receptors, we further varied the number of antibodies in the simulations.

**Figure 2:**
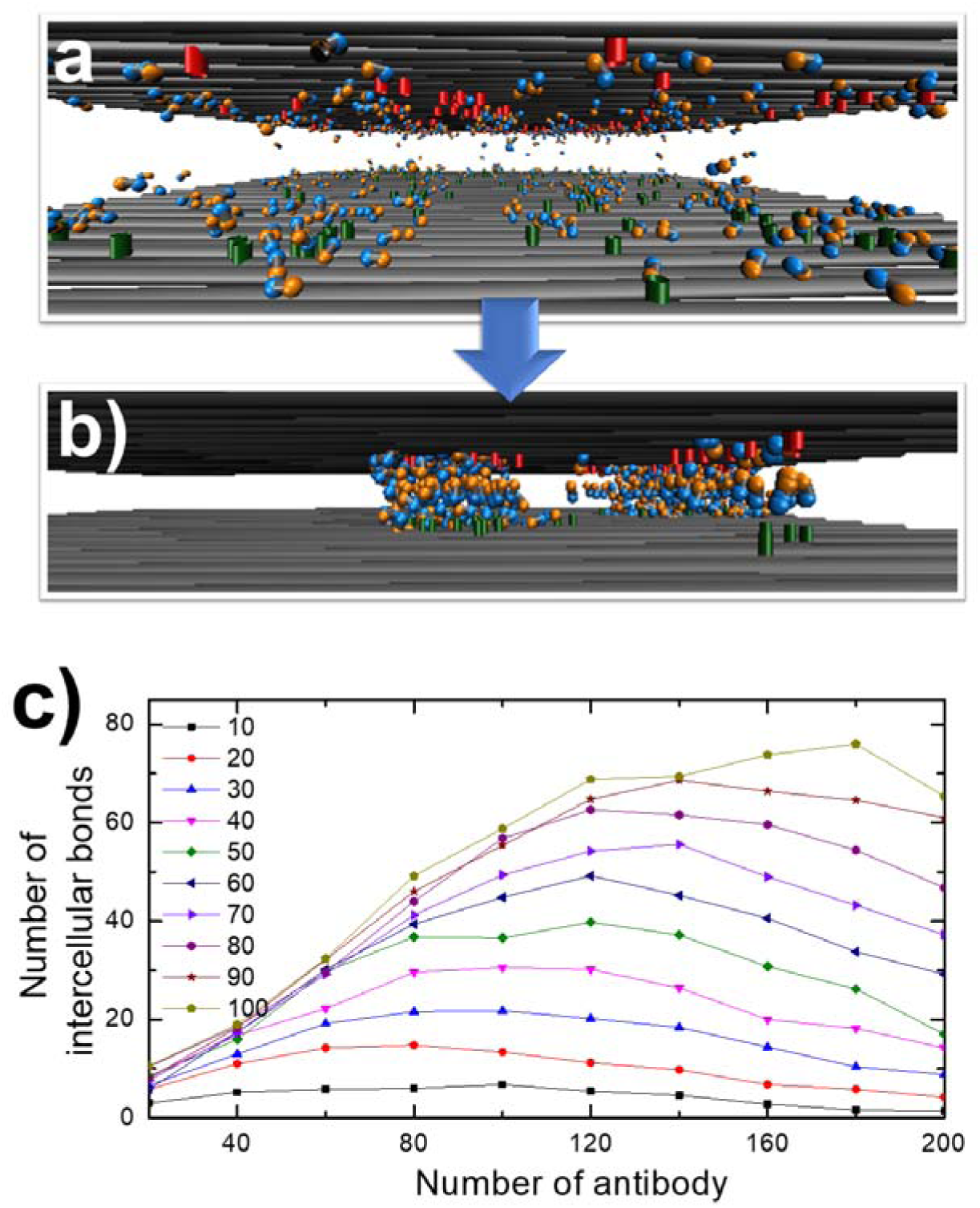
The side views of the initial and final configurations from a representative trajectory of the intercellular simulation are shown in **(a)** and **(b)**, respectively. The plasma membranes are represented by the gray surfaces. Receptors CD3 and TAA are represented by the red and green cylinders. The two functional modules of a bispecific antibody are represented by orange and blue spheres and their linker is shown in gray. Intercellular simulations were carried out for different numbers of antibody and receptor. The changes of intercellular bonds averaged at the end of five independent trajectories were plotted in **(c)** as a function of antibody number. Each curve in the figure represents a specific value of receptor density, which is indexed on the left side.

Given different numbers of antibody and receptor in the system, five independent simulation trajectories were carried out. At the end of each trajectory, the number of intercellular bonds, i.e., the number of ternary complexes formed among CD3, antibody and TAA, was calculated. To examine the correlation between receptor density and antibody concentration, the binding affinity between antibodies and receptors in all simulations was fixed at a very strong level (−21kT), corresponding to the dissociation constant of ∼0.1nM. **Figure 2c** shows the average number of intercellular bonds over all trajectories under different conditions. Each curve in the figure represents a specific value of receptor density and shows the change in the average number of intercellular bonds as a function of antibody concentration. For all values of receptor surface density, a concave down dependence on antibody concentration was observed. Specifically, at low antibody concentrations, the number of intercellular bonds increased with increasing antibody concentration, reaching a maximum number and then decreasing with continued increase in antibody concentration. This change of intercellular bonds as a function of antibody concentration is known as the classic Hook effect, which is anticipated in the theory of three-body binding equilibria, and which has been observed in recent experiments on bispecific T cell engagers [42].

The mechanistic basis for the decrease in intercellular bonds at high antibody concentrations is the consequence of high and saturated levels of binary complexes on both cell surfaces, which preclude the formation of ternary complexes. Furthermore, as the surface density of receptors increased, we discovered that not only did the peak’s height increase, but its position also shifted from left to right. This behavior suggests that a higher surface density of receptors can result in stronger adhesion between cells (more crosslinking interactions) and, at the same time, require a higher concentration of antibodies to observe the Hook effect, both of which may be important clinical considerations.

We next investigated the sensitivity of the Hook effect to the affinity between antibodies and receptors. We examined five different binding affinity, spanning a wide range from a very weak interaction of -5kT (corresponding to an equilibrium dissociation constant of 10 mM) to a very strong interaction of -21kT (corresponding to an equilibrium dissociation constant of 0.1 nM). In these simulations, the number of receptors on both surfaces were fixed at 100. As in previous experiments, the size of each square surface was 500nm along both the x and y directions, and the distance between two surfaces was reduced from 70nm to 35nm along the simulations. For each value of binding affinity, five independent simulation trajectories were performed employing different numbers of antibody and the average number of intercellular bonds computed.

**Figure 3a** displays the average number of intercellular bonds as a function of antibody concentration for different binding affinities. The Hook effect was observed in all systems. However, the positions of the maxima were dependent on the values of binding affinity. The correlation between the peak position and the corresponding binding affinity is displayed in **Figure 3b** and shows that higher concentrations of low affinity antibodies are required to observe the Hook effect. When the interaction between antibodies and receptors is strong, binary complexes are relatively easier to form. As a result, a small number of antibodies can occupy the majority of receptors on both cell surfaces, preventing them from forming ternary complexes. On the other hand, when the interaction between antibodies and receptors is weak, binary complexes are relatively more difficult to form. As a result, the system needs a larger number of antibodies to prevent receptors from forming ternary complexes.

**Figure 3:**
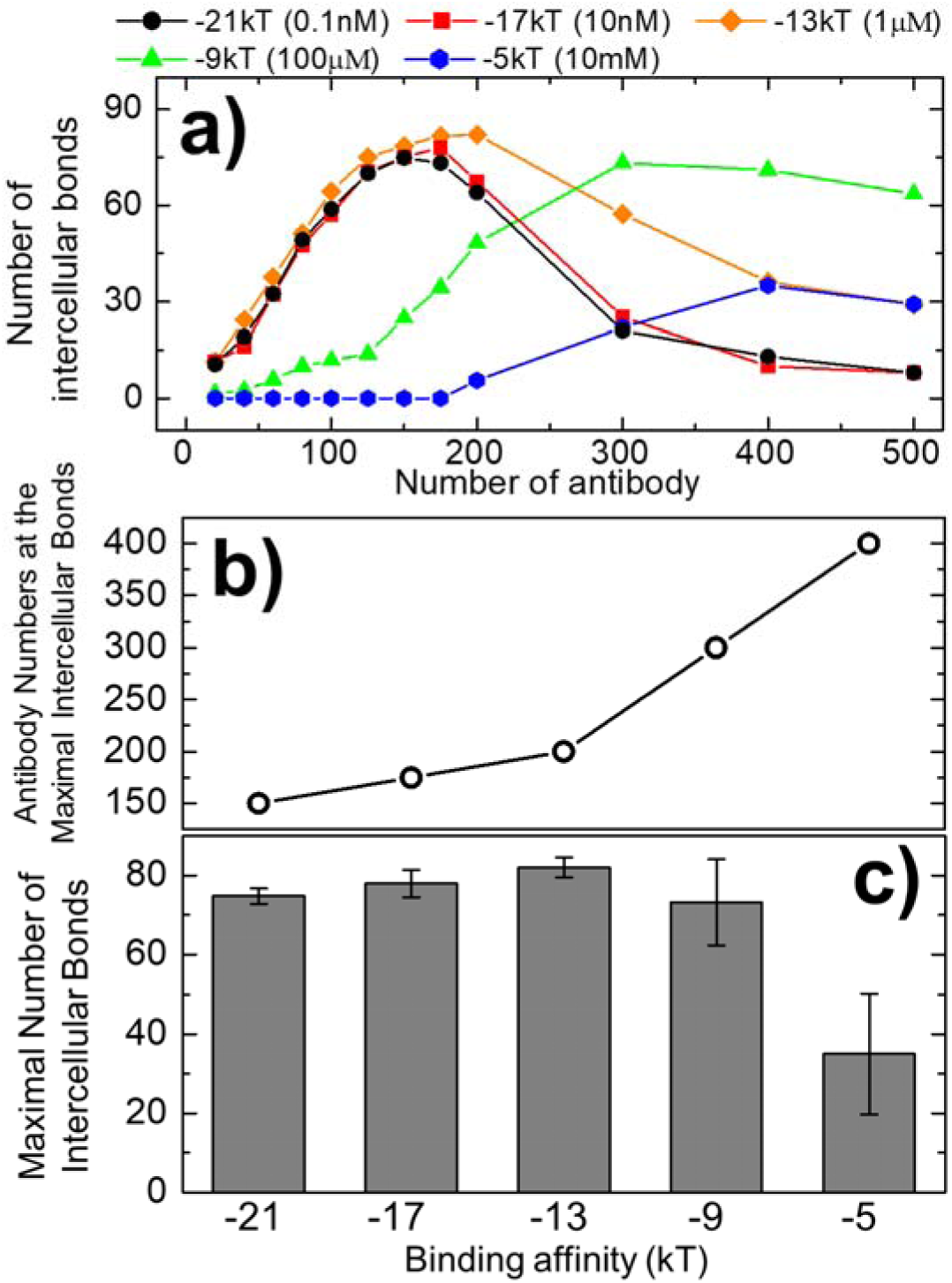
We tested five different binding affinities between receptors and antibodies. Their changes of average intercellular bonds are shown in **(a)** under different antibody concentrations. The symbols of binding affinities in the figure are listed on the top. A peak was observed in each system. The correlation between the peak position, i.e., the antibody concentration when the intercellular bonds reach the maximal value, and the corresponding binding affinity was plotted in **(b)**. The heights of the peaks, i.e., the maximal value of average intercellular bonds, under different binding affinities were further plotted in (c).

Moreover, we found that the height of the peaks, i.e., the maximal value of average intercellular bonds, also changed with the antibody concentration (**Figure 3c**). Interestingly, the maximal values of intercellular bonds do not change monotonically with the binding affinities. Intuitively, the number of intercellular bonds a system can achieve might be expected to decrease as the binding affinity between receptors and antibodies in the system weakens. However, **Figure 3c** indicates that when the binding affinities became weaker, the heights of the corresponding peaks increased first, reached the maximal value, and then decreased. We thus speculate that the formation of intercellular bonds is not only determined by the binding affinity between receptors and antibodies, but can also be affected by the spatial pattern of the ternary complexes at the cell-cell interface.

To quantify the spatial organization, we visualized the positions of all receptors and antibodies generated at the end of the simulations. We found that the CD3 and TAAs can cluster together through their binding to the bispecific antibodies. We manually determined the size of each cluster, and the maximal cluster was selected for each trajectory, and their average value was calculated from all five trajectories. These average values of the maximal clusters were plotted as histograms with error bars in **Figure 4a** under different binding affinities. If the binding between antibodies and receptors is as weak as -5kT (10 mM in dissociation constant), only small clusters were observed due to the low number of intercellular bonds. Under stronger binding affinities, the simulations suggested that the maximal size of the clusters formed in the system had a negative correlation with the binding affinity. For instance, the clusters formed under the binding affinity of -21kT are much smaller than the clusters formed under the binding affinity of - 9kT.

**Figure 4:**
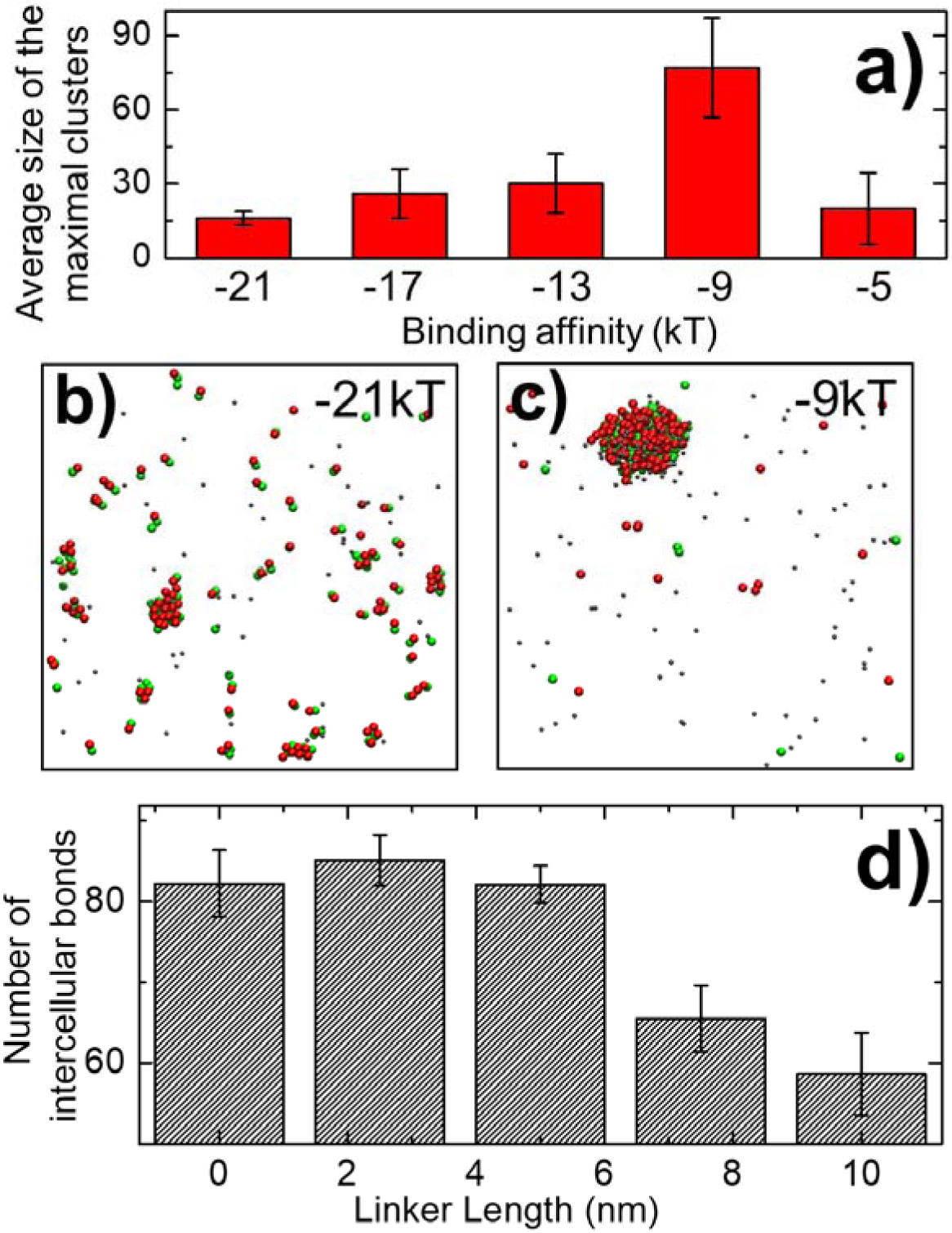
Under each different value of binding affinity, we calculated the average size of the maximal clusters formed at the end of all five trajectories **(a)**. Some representative snapshots were selected from the end the simulations. The top views of the cell interface under the binding affinity of -21kT and -9kT are shown in **(b)** and **(c)**, respectively. The CD3, TAAs, and antibodies are represented by the red, green and gray dots in the figures. We also carried out simulations by changing the length of the linkers in the bispecific antibodies. The average numbers of intercellular bonds under different linker lengths are summarized in **(d)** as histograms.

Representative snapshots were selected from the end of the simulations of these two binding affinities. The top view of the cell-cell interface observed with the binding affinity of -21kT is shown in **Figure 4b**, while the top view of the cell-cell interface observed with the binding affinity of -9kT is shown in **Figure 4c**. The CD3 receptors on the top cell surface are shown by the red dots in the figures, the TAA receptors on the bottom cell surface are shown by the green dots and the bispecific antibodies are shown by the grey dots. In both snapshots, an intercellular bond is represented by a colocalized pair of red and green dots. These colocalized intercellular receptors further aggregated together into different clusters. When the binding affinity equals -21kT, almost all CD3 and TAA formed intercellular bonds. These ternary complexes further aggregated into several small clusters. In contrast, when the binding affinity equals -9kT, although considerable amounts of receptors were not involved in the intercellular bonds, a single large cluster was formed. These are consistent to the histogram in **Figure 4a**.

Antibody binding slows the diffusions of receptors on cell surface due to the increased mass of the complexes. These antibody-receptor complexes serve as seeds to attract nearby unoccupied receptors through non-specific collisions. The newly joined receptors then further bind to antibodies, and in turn the receptors on the other side of the cell surface. Through this feedback process, a cluster can be formed, and its size can keep growing. Under strong affinity, receptors and antibodies have low probability to dissociate. The ternary complexes are then kinetically trapped in small clusters. In contrast, when the affinity is reduced, the clustering process is more dynamic. Clusters are more frequently dissolved and merged with each other, leading to the final formation of few but larger clusters. Moreover, the ternary complexes formed in the middle of a larger cluster are more difficult to dissociate due to the higher local concentration, which explains the higher number of intercellular bonds under relatively weaker binding affinities. The clustering of receptors plays an important role in regulating the downstream cell signaling pathways. The above studies therefore suggest that the highest affinity binding may not result in the optimal therapeutic outcome. Interestingly, our simulation results provide a mechanistic explanation to a recent study which showed the intermediate affinity, instead of high, affinity of antibodies can lead to a higher level of signaling activity through increased receptor clustering [43].

The two functional modules in a bispecific antibody are connected by a linker. Previous studies showed that the binding of a fusion protein to its targeting receptors can be modulated by the physical properties of its linkers [44]. To examine the impact of linker length within the bispecific antibody on T cell engagement, we performed simulations with different length linkers. Specifically, five systems were prepared, in which the linker length was gradually increased from 0 nm to 10 nm. The final inter-membrane distance was accordingly adjusted corresponding to the linker length in each system. For instance, if the linker length equals 5nm, the intermembrane distance was reduced from 70nm to 35nm along the simulation, as described above. Similarly, if the linker length equals 0nm, the intermembrane distance was reduced to 30nm. On the other hand, if the linker length equals 10nm, the intermembrane distance was reduced to 40nm. All other parameters in these five systems were the same: the binding affinity was fixed at 13kT; there were 100 receptors on each cell surface and 200 antibodies in the intercellular space; the size of the cell surface is 500nm×500nm in area. For each system, five independent simulation trajectories were generated and the average number of intercellular bonds were calculated after all simulation completed.

The results of average intercellular bonds under different linker lengths are summarized in **Figure 4d** as histograms with error bars. The figure shows that more intercellular bonds were formed as the linker length increased from 0 nm to 2.5 nm, after which fewer intercellular bonds formed as the linker length further increased to 10 nm. This result suggests that there is an ideal length for the linker of the bispecific antibody to optimize its effect on T cell engagement. If the linker is too short, we speculate that the local searching space of an antibody for its targeting receptor will be negatively impacted, so that fewer ternary complexes are formed. On the other hand, if the linker is too long, we speculate that the increased amplitude in orientational degrees of freedom in the antibodies make their binding to cell surface receptors more difficult, and thus prevent them from forming more ternary complexes. As a result, our simulations provide insights to the molecular design of bispecific T cell engager with optimal linker length to maximize the adhesion strength between T cells and tumor cells.

The results of the intercellular simulations were used to generate the input for multicellular simulations of the T cell engaging process. We compared the interaction between a T cell and a tumor cell to the interaction between a T cell and a normal cell, as shown in **Figure 5a**. We placed 100 CD3 molecules on the surface of a T cell and 100 TAAs on the surface of a tumor cell on the opposite side (surface dimensions of 500nm×500nm). Because it is possible that TAAs could also be expressed in normal cells, three different expression levels were tested. For the low expression level, 10 TAAs were placed on the surface of a normal cell. For the medium and high expression levels, 20 and 50 TAAs were included in the systems, respectively. For each intercellular system, we further changed the antibody-receptor binding affinity and antibody concentration, to test how the multicellular interactions respond to these factors. Specifically, two binding affinities, -9kT and -13kT, were tested, and 50, 200 and 500 antibodies per surface area were used to represent the low, medium and high concentrations. Five independent trajectories of the intercellular simulations were performed for each of these conditions. The **average number** of intercellular bonds was calculated for each condition. As described in the methods, given the size of interface area, these averages of the number of intercellular bonds were used as parameters to model the adhesion strengths between T cells, tumor cells and normal cells in the multicellular simulations.

**Figure 5:**
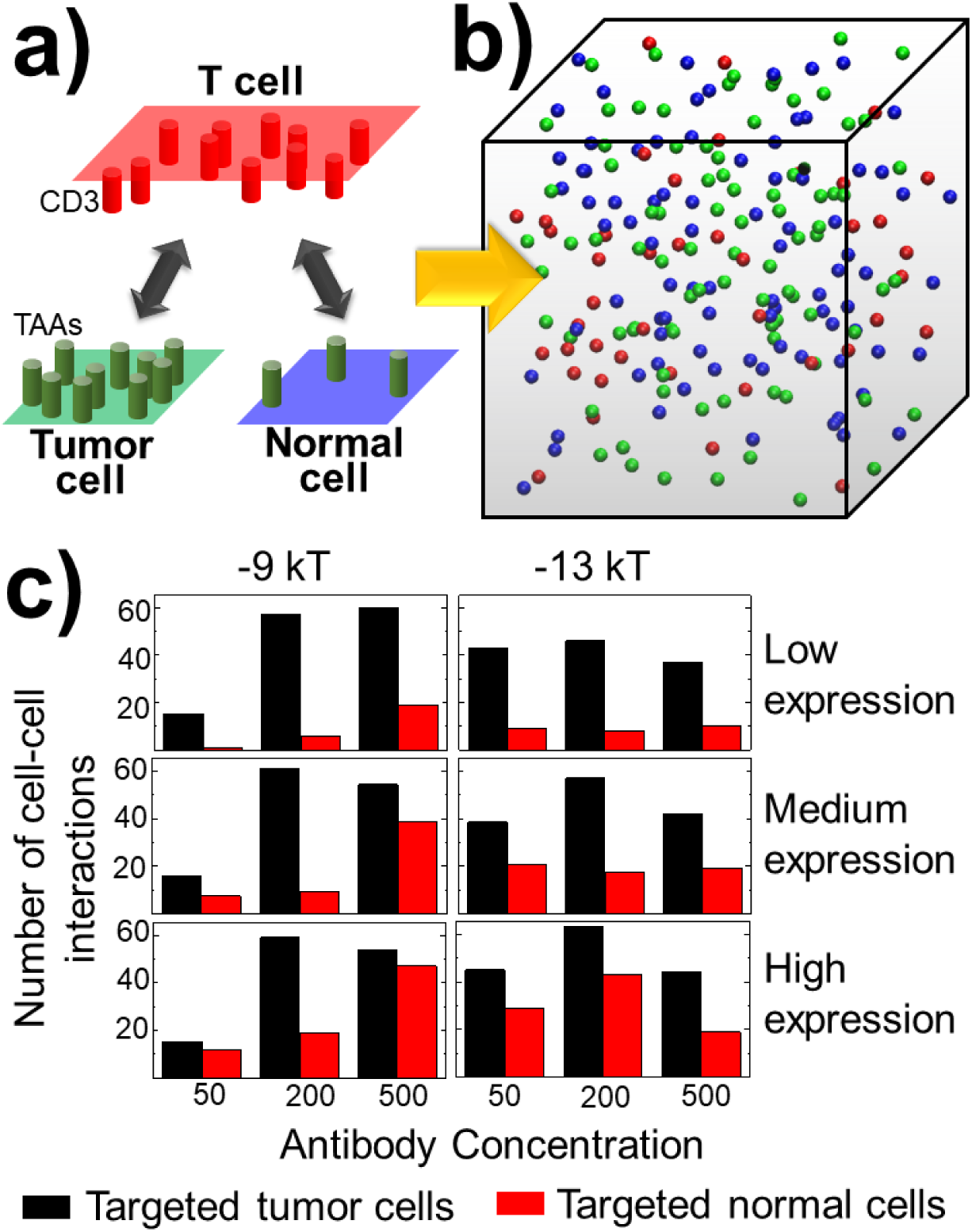
The intercellular simulations were used to generate the input for the multicellular simulations of the T cell engaging process **(a)**. We first derived the number of intercellular bonds formed between a T cell and a tumor cell, and the number of intercellular bonds formed between a T cell and a normal cell under various combinations of TAA expression levels, antibody concentrations, and binding affinities. The results were then fed into a multicellular system, of which the initial configuration is shown in **(b)**. After the simulations, we calculated how many tumor cells were targeted by the T cells, and compared them to the number of normal cells that were also targeted by the T cells. The results were summarized in **(c)** under all different conditions.

In the multicellular simulations, initial configurations were generated by randomly distributing 200 cells of radius 5μm in a simulation box, of dimension 240×240×240 μm^3^. The cells were selected from among three types in the fixed ratio of 40 of T cells (red in **Figure 5b**), 80 tumor cells (green in **Figure 5b**) and 80 normal cells (blue in **Figure 5b**). After the simulations, we calculated the number of tumor cells targeted by the T cells, and compared them to the number of normal cells targeted by the T cells. Results are summarized in **Figure 5c**, with the number of tumor and normal cells targeted by the T cells represented by the black and red bars, respectively. The left panels denote the systems in which the antibody-receptor binding affinity equals -9kT, while the right panels denote the systems in which the antibody-receptor binding affinity equals - 13kT. The systems in which the normal cells contain the low expression level of TAAs were plotted by the top two panels. The systems in which the normal cells contain medium and high expression levels of TAAs were plotted by the medium and bottom two panels, respectively. Each panel of **Figure 5c** contains three groups of data, corresponding to the systems with the low, medium and high antibody numbers that are represented by the black and red bars on the left, medium and right.

**Figure 5c** shows that if a specific TAA has a low expression level in the normal cells, the number of targeted tumor cells is much higher than the number of target normal cells under all binding affinities and antibody concentrations. When the expression level of TAA increased in the normal cells, the differences between the number of targeted tumor cells and targeted normal cells became smaller. Under medium and high TAA expression levels, we found that the number of targeted tumor cells was reduced if the number of antibodies in the systems was too high, as shown by the black bars on the right side of each panel in the middle and bottom rows. This result suggests that the high antibody concentration might negatively affect the efficiency of T cell engagement due to the Hook effect described before. Moreover, by comparing the simulation results between two binding affinities, we suggest that the performance of the stronger affinity antibody is more stable. The numbers of targeted tumor cells are always reasonably higher than the numbers of targeted normal cells under all conditions, as reflected by the right three panels in **Figure 5c**. In contrast, when the binding affinity is weaker, the difference between the number of targeted tumor cells and normal cells became marginal under low and high antibody concentrations, as reflected by the left three panels. In summary, these multicellular simulations under various combinations of molecular and cellular determinants gave us an overview of how to adopt the most appropriate strategy to maximize the drug efficacy and avoid the off-target effect to the greatest extent.

There is no adhesion between tumor cells and no adhesion between normal cells in the above multicellular simulations, which may be more representative of liquid tumors. However, in most of the solid tumors, these cells can form interactions with each other. In order to understand the impact of cell adhesion on T cell engagement, we included interactions between tumor cells and the interactions between normal cells into our multicellular simulations. Specifically, two simulation scenarios were compared with each other. In the first scenario, there were no adhesive interaction between tumor cells, nor between normal cells; this the tumor and normal cells can only interact with T cells. In the second scenario, adhesive interactions between tumor cells and between normal cells were added to the simulation besides their interactions with T cells. Other simulation parameters in these two systems were the same. In both simulations, 125 cells were randomly distributed in a cubic simulation box as an initial configuration. 25 of them are T cell; 50 of them are tumor cells; and the rest 50 of them are normal cells. The volume of the simulation box is 200×200×200 μm^3^. The snapshots of the final configurations from the simulations of the first and the second scenario are shown in **Figure 6a** and **Figure 6b**, respectively. The red, green and blue spheres in the figures represent T cells, tumor cells and normal cells.

**Figure 6:**
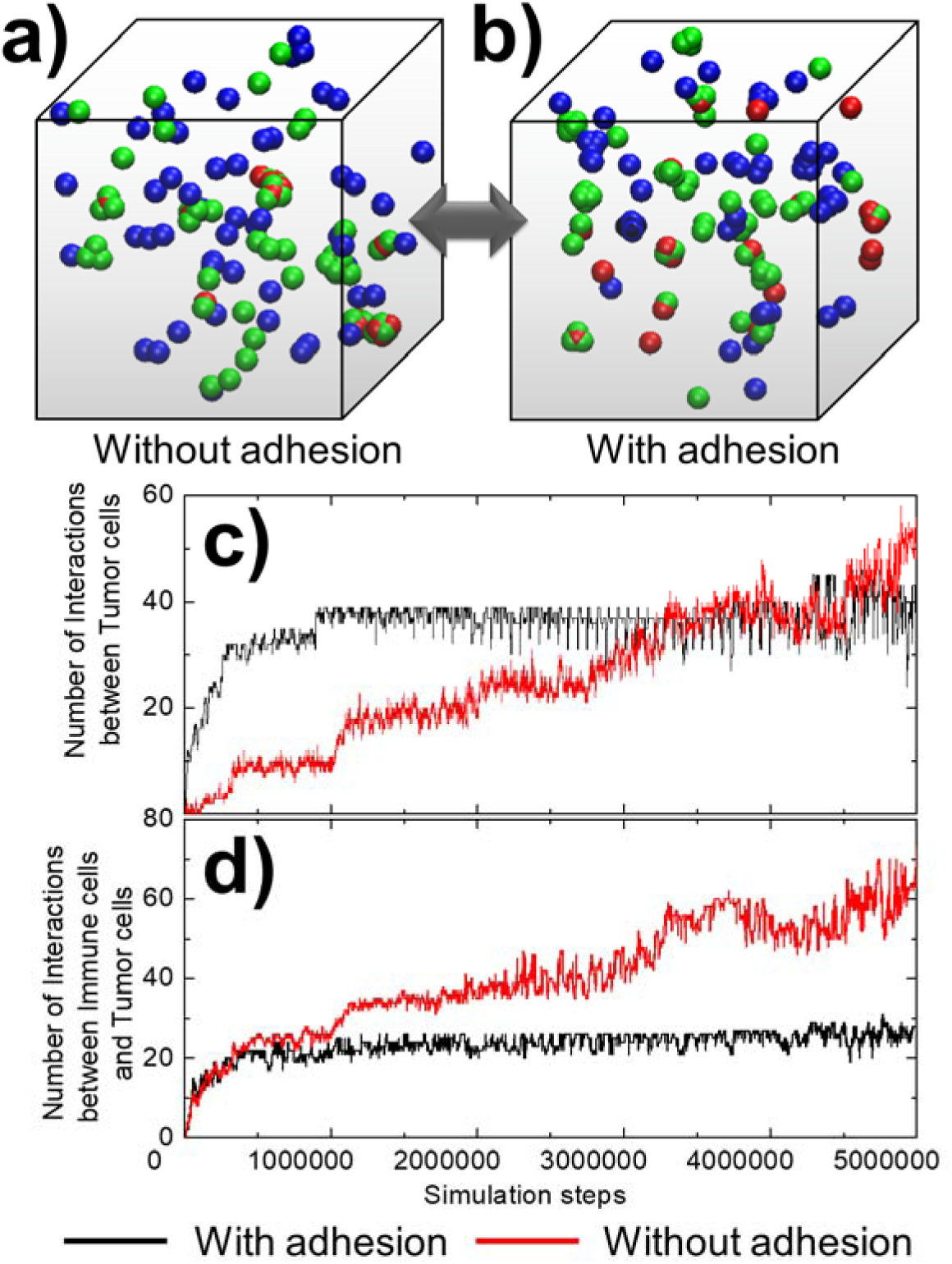
We introduced the interactions between tumor cells and the interactions between normal cells into our multicellular simulations. The snapshots of the final configurations from the simulations without these interactions and the simulations with these interactions are shown in **(a)** and **(b)**, respectively. The kinetic profiles of the simulations as a function of time are further compared between these two systems. In detail, the total numbers of interactions between tumor cells formed along the simulation were plotted in **(c)**, while the total numbers of interactions formed between T cells and tumor cells along the simulation were plotted in **(d)**. The results from the simulations without adhesion between tumor cells and adhesion between normal cells are represented by the red curves in both figures, and the results from the simulations with adhesion between tumor cells and adhesion between normal cells are represented by the black curves.

The kinetic profiles of the simulations as a function of time gave a more quantitative comparison of the two systems. In detail, the total numbers of interactions between tumor cells formed along the simulation were plotted in **Figure 6c**, while the total numbers of interactions formed between T cells and tumor cells along the simulation were plotted in **Figure 6d**. The red curves in the figures represent the kinetic profiles of the first simulation scenario in which no adhesion existed between tumor cells and between normal cells. The black curves represent the kinetic profiles of the second simulation scenario in which adhesion was introduced between tumor cells and between normal cells. **Figure 6b** shows that small clusters of tumor cells and of normal cells were formed after introducing adhesive interactions. As a result, the system reached equilibrium at an early stage of the simulation, when the number of interactions between T cells and tumor cells stopped growing at a relatively low level. On the other hand, when there is no adhesion between tumor cells and between normal cells, the number of interactions between T cells and tumor cells kept growing to a much higher level through the end of the simulation. Moreover, **Figure 6a** shows that a T cell can simultaneously interact with multiple tumor cells. This is also confirmed by the red curve in **Figure 6c**, in which the number of interactions between tumor cells was very low at the beginning, but increased to a high level at the end of the simulation. In summary, our results suggest that the adhesion between tumor cells can prevent their engagements with T cells. This also explains why T cell engagers are much less effective in the treatment of solid tumors than liquid tumors [45].

## Concluding Discussions

Under physiological conditions, only when the T-cell receptors (TCR) on the surface of a T cell recognize their corresponding epitopes can the cytotoxic activity of the T cell be directed towards the cells expressing major histocompatibility class (MHC) molecules that present these epitopes [46]. Cancer cells have developed different strategies to escape immune surveillance, many of which are related to how MHC molecules process and present antigens [47]. By directly linking CD3 and TAA through T-cell engagers, the T-cell activation triggered by the normal TCR-MHC interaction can be bypassed. The applications of these bispecific antibodies to redirect the cytotoxic activity of T-cells in a non-MHC restricted fashion have thus drawn enormous attention in recent years. However, the detailed mechanisms by which bispecific antibodies regulate the physical adhesion between T cells and tumor cells have not been well studied. Understanding this process on a more quantitative level can help to overcome the challenges commonly experienced in clinical applications of T cell engagers, including on-target off-tumor toxicity, tumor antigen escape and suboptimal potency.

To achieve these goals, we developed computational methods to study the process of T cell engagement on the intercellular and multicellular scales. On the intercellular level, kinetic Monte-Carlo simulation was first used to study the molecular determinants (e.g., CD3 and TAA) that affect the intercellular bonds formed by bispecific antibodies. We found that the correlation between the number of intercellular bonds and the antibody concentration follows the classic Hook effect. Specifically, if the antibody concentration is higher than a threshold, a majority of receptors will be occupied by excess numbers of antibodies. These binary complexes will prevent the further formation of intercellular ternary complexes. We showed a sensitive relationship between the threshold of the Hook effect and the maximal number of intercellular bonds. We found that the ternary complexes can aggregate into clusters at the cell-cell interfaces, and that the size of the clusters depends on the binding affinity between receptors and antibodies. More interestingly, our results suggest that the medium-level binding affinity between antibodies and receptors can lead to the formation of larger clusters than the strong binding affinity. Finally, an optimal value in the length of the linker was observed in the bispecific antibody.

On the multicellular level, the Langevin dynamics simulation was used to drive the spatial-temporal dynamics of adhesions among T cells, tumor cells and normal cells. The strengths of adhesions between different types of cells were transferred from the number of intercellular bonds formed between CD3 and TAA that were derived by the previous intercellular simulations. By testing different combinations of TAA expression levels, receptor binding affinities and antibody concentrations, we showed that the high antibody concentrations negatively affected the efficiency of T cell engagement and the antibody with stronger affinity was more stable in directing tumor cells to T cells. Furthermore, demonstrated that adhesive interactions between tumor cells and between normal cells can impair their engagement with T cells, which explains why T cell engagers are more effective in the treatment of liquid tumors than solid tumors. The current multiscale framework, which integrates the intercellular and multicellular simulations, will serve as a computational platform to help the future design of new biological therapeutics.

As a proof-of-concept study, our current simulation methods include a few simplifications. Further improvements will be made to address these limitations. First, the plasma membrane of cells is simplified as a flat surface in current study by assuming that the major impact of membrane environments on the dynamics of cell surface proteins is to constrain their diffusions inside of the plasma membrane. However, membrane undulations might also play a role in regulating the interactions between antibodies and cell surface receptors [48]. We will model the plasma membrane as an elastic medium by following the procedure described in our recent study. Second, both our intercellular and multicellular simulations start from a random initial configuration. In living cells, however, the distributions of membrane receptors on the plasma membrane are not random, but rather highly heterogeneous. The information of this distribution can be derived from available experimental evidences such as the data reported from super-resolution imaging [49]. Similar strategy could be applied to obtain the spatial organization of T cells and tumor cells in a specific tissue environment. Finally, the linker flexibility in a bispecific antibody was neglected in current simulation. Our recent study revealed that both length and flexibility of the linker in a multidomain protein ligand can change its conformational dynamics [50] and further affect the binding to its targeting receptor. The conformational variations of a bispecific antibody will be incorporated into our intercellular simulations in the future.

## Acknowledgement

This work was supported by the National Institutes of Health under Grant Numbers R01GM120238 and R01GM122804 (to YW), R01AI145024 and R01CA198095 (to SCA). We acknowledge the Wollowick Family Foundation Chair in Multiple Sclerosis and Immunology (to SCA), and Janet & Martin Spatz and the Helen & Irving Spatz Foundation. We also acknowledge the Einstein-Rockefeller-CUNY Center for AIDS Research (P30AI124414) and the Albert Einstein Cancer Center (P30CA013330). The work is also partially supported by a start-up grant from Albert Einstein College of Medicine. Computational support was provided by Albert Einstein College of Medicine High Performance Computing Center.

## Author Contributions

Z.S., S.C.A. and Y.W. designed research; Z. S. and Y.W. performed research; Z.S., and Y.W. analyzed data; S.C.A. and Y.W. wrote the paper.

## Additional Information

**Competing financial interests:** The authors declare no competing financial interests.

